# Real-time detection of PRT1-mediated ubiquitination via fluorescently labeled substrate probes

**DOI:** 10.1101/062067

**Authors:** Augustin C. Mot, Erik Prell, Maria Klecker, Christin Naumann, Frederik Faden, Bernhard Westermann, Nico Dissmeyer

## Abstract

- The N-end rule pathway has emerged as a major system for regulating protein functions by controlling their turn-over in medical, animal and plant sciences as well as agriculture. Although novel functions and enzymes of the pathway were discovered, ubiquitination mechanism and substrate specificity of N-end rule pathway E3 Ubiquitin ligases remained elusive. Taking the first discovered *bona fide* plant N-end rule E3 ligase PROTEOLYSIS1 (PRT1) as a model, we use a novel tool to molecularly characterize polyubiquitination live, in real-time.
- We gained mechanistic insights in PRT1 substrate preference and activation by monitoring live ubiquitination by using a fluorescent chemical probe coupled to artificial substrate reporters. Ubiquitination was measured by rapid in-gel fluorescence scanning as well as in real time by fluorescence polarization.
- Enzymatic activity, substrate specificity, mechanisms and reaction optimization of PRT1-mediated ubiquitination were investigated *ad hoc* in short time and with significantly reduced reagent consumption.
- We demonstrated for the first time that PRT1 is indeed an E3 ligase, which was hypothesized for over two decades. These results demonstrate that PRT1 has the potential to be involved in polyubiquitination of various substrates and therefore pave the way to understanding recently discovered phenotypes of *prt1* mutants.

## INTRODUCTION

The ON/OFF status of protein function within the cells’ proteome, their general abundance and specific distribution throughout the compartments and therefore their functions and activities are precisely controlled by protein quality control (PQC) mechanisms to ensure proper life of any organism. Therefore, the biochemical analysis of the underlying mechanisms safeguarding proteostatic control is pivotal. It ranges from the molecular characterization of enzymes involved in PQC and their catalyzed reactions to enzyme-substrate and non-substrate protein-protein interactions. The so-called Ubiquitin (Ub) 26S proteasome system (UPS) is a master component of PQC with the key elements being non-catalytic Ub ligases (E3), the Ub-conjugating enzymes (E2), and the Ub-activating enzymes (E1).

To investigate an element conferring substrate specificity, we chose PROTEOLYSIS1 (PRT1) from *Arabidopsis thaliana* as a model E3 ligase, which is a *bona fide* single-subunit E3 with unknown substrate portfolio (Bachmair *et al.*, 1993; Potuschak *et al.*, 1998; Stary *et al.*, 2003). Its biological function remains elusive but it presumably represents a highly specific enzyme with E3 ligase function of the N-end rule pathway of targeted protein degradation, which is a part of the UPS. The N-end rule relates the half-life of a protein to its N-terminal amino acid (Bachmair *et al.*, 1986) and causes rapid proteolysis of proteins bearing so-called N-degrons, N-terminal sequences that lead to the degradation of the protein. N-degrons are created by endoproteolytic cleavage of protein precursors (pro-proteins) and represent the resulting neo-N-termini of the remaining C-terminal protein moiety, albeit not all freshly formed N-termini automatically present destabilizing residues (**Fig. 1a**).

**Figure 1.**
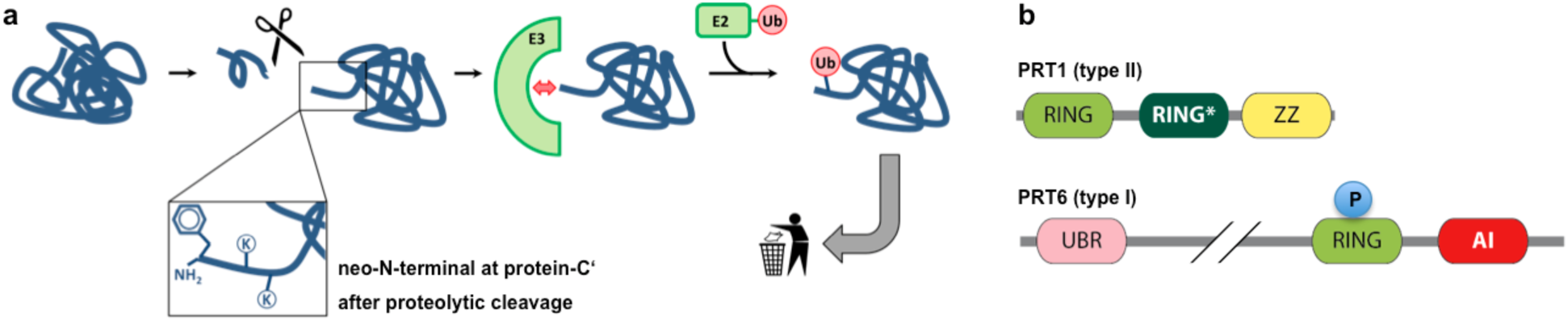
Generation of N-end rule substrates by proteolytic processing and predicted features of the two *bona fide* plant N-recognins. **a)** Substrates containing N-degrons can be generated from (pre-)pro-proteins as precursor sequences after proteolytic cleavage (indicated by the scissors). The N-degron shown here comprises a Phe residue as primary destabilizing residue at the protein-C’ and internal lysines for polyubiquitination. These N-degrons can be recognized by N-end rule E3 Ub ligases (N-recognins) which in turn associate with Ub-conjugating enzymes (E2) carrying Ub which was previously activated by E1 enzymes. One possible result of ubiquitination is protein degradation and to date, in the context of the N-end rule, ubiquitination is assumed to lead to degradation in most of the cases. **b)** The two known *Arabidopsis* N-recognins were identified by their function (PRT1, 46 kDa) and by homology to the UBR-box from*S. cerevisiae* UBR1p (PRT6, 224 kDa). UBR: box binding type I substrates; RING*: composite domain containing RING and CCCH-type Zn fingers; ZZ: Zinc binding domain similar to RING; RING: protein-protein interaction domain for E2–E3 interaction; AI: predicted autoinhibitory domain (intramolecular interaction); P: phosphorylation site (PhosPhAt 4.0; phosphat.unihohenheim.de). b is modified from Tasaki et al., 2012.

The N-end rule pathway is an emerging vibrant area of research and has a multitude of functions in all kingdoms (Dougan *et al.*, 2010; Varshavsky, 2011; Tasaki *et al.*, 2012; Gibbs *et al.*, 2014a; Gibbs, 2015). Identified substrates are mainly important regulatory proteins and play key roles in animal and human health (Zenker *et al.*, 2005; Piatkov *et al.*, 2012; Brower *et al.*, 2013; Shemorry *et al.*, 2013; Kim *et al.*, 2014), plant stress response and agriculture (Gibbs *et al.*, 2011; Licausi *et al.*, 2011; Gibbs *et al.*, 2014a; Gibbs *et al.*, 2014b; Weits *et al.*, 2014; de Marchi *et al.*, 2016; Mendiondo *et al.*, 2016).

In plants, functions of N-end rule enzymes are associated with central developmental processes including seed ripening and lipid breakdown, hormonal signaling of abscisic acid (ABA), gibberellin and ethylene, seed dormancy and germination (Holman *et al.*, 2009; Abbas *et al.*, 2015; Gibbs *et al.*, 2015), leaf and shoot morphogenesis, flower induction, and apical dominance (Graciet *et al.*, 2009), and the control of leaf senescence (Yoshida *et al.*, 2002). Then, the pathway was shown to be a sensor for molecular oxygen and reactive oxygen species (ROS) by mediating nitric oxide (NO) signaling and regulating stress response after hypoxia, e.g. after flooding and plant submergence (Gibbs *et al.*, 2011; Licausi *et al.*, 2011; Gibbs *et al.*, 2014b). A novel plant-specific class of enzymes was associated with the pathway, i.e. plant cysteine oxidases (PCOs), highlighting plant-specific molecular circuits, enzyme classes and mechanisms (Weits *et al.*, 2014). In the moss *Physcomitrella patens*, N-end rule mutants are defective in gametophytic development (Schuessele *et al.*, 2016) and protein targets of N-end rule-mediated posttranslational modifications were discovered (Hoernstein *et al.*, 2016). Also in barley, the pathway is connected with development and stress responses (Mendiondo *et al.*, 2016). Only very recently, a link between N-end rule function and plant-pathogen response and innate immunity was found (de Marchi *et al.*, 2016), shedding light on novel functions of the yet underexplored branch of targeted proteolysis. However, to date, the identity of plant N-end rule targets still remains obscure and clear evidences from biochemical data of *in vitro* and *in vivo* studies such as N-terminal sub-proteomics or enzymatic assays are still lacking.

A novel *in vivo* protein stabilization tool for genetic studies in developmental biology and biotechnological applications, the 'lt-degron', works in plants and animals by directly switching the levels of functional proteins *in vivo* (Faden *et al.*, 2016). The method is based on conditional and specific PRT1-mediated protein degradation, the process studied in depth with the here-generated fluorescent substrate reporters.

N-degrons are by definition recognized and the corresponding protein ubiquitinated by specialized N-end rule E3 ligases, so-called *N-recognins* (Sriram *et al.*, 2011; Varshavsky, 2011; Tasaki *et al.*, 2012; Gibbs, 2015). In plants, only two of these, namely PRT1 and PRT6, are associated with the N-end rule and assumed to function as N-recognins (**Fig. 1b**). This is in contrast to the high number of proteolytically processed proteins which carry in their mature form N-terminal amino acids that could potentially enter the enzymatic N-end rule pathway cascade (Venne *et al.*, 2015). In the light of more than 800 putative proteases in the model plant *Arabidopsis thaliana*, it is likely that the N-end rule pathway plays an important role for protein half-lives in a proteome-wide manner. Examples are found in the METACASPASE9 degradome, i.e. that part of the proteome which is associated with degradation (Tsiatsiani *et al.*, 2013), or the N-degradome of *E. coli* (Humbard *et al.*, 2013) with a possibly analogous overlap with endosymbiotic plant organelles (Apel *et al.*, 2010).

PRT1, compared to the *Saccharomyces cerevisiae* N-recognin Ubr1 (225 kDa), is a relatively small protein (46 kDa) and totally unrelated to any known eukaryotic N-recognins but with functional similarities to prokaryotic homologs (**Fig. 1b**). It is therefore perceived as a plant pioneer E3 ligase with both diversified mechanistics and function. Artificial substrate reporters based on mouse dihydrofolate reductase (DHFR) comprising an N-terminal phenylalanine generated via the ubiquitin-fusion (UFT) technique lead to the isolation of a *prt1* mutant in a forward mutagenesis screen (Bachmair *et al.*, 1993). In the mutant cells and after MG132 treatment, the F-DHFR reporter construct was shown to be stabilized whereas it was instable in the untreated wild type (Potuschak *et al.*, 1998; Stary *et al.*, 2003). *PRT1* was able to heterologously complement a*Saccharomyces cerevisiae ubr1Δ* mutant strain where Phe-, Tyr-, and Trp-initiated β-galactosidase test proteins were stabilized. These reporters were rapidly degraded in *ubr1Δ* transformed with PRT1 (Stary *et al.*, 2003). A new study revealed that cleavage of the E3 ligase BIG BROTHER by protease DA1 forms a C-terminal, Tyr-initiated fragment. Its stability depends on the N-terminal amino acid Tyr and the function of PRT1 E3 ligase (Dong *et al.*, 2016). However, until today, there are no more *in vivo* targets or direct functions associated with PRT1, but recently, a potential role of PRT1 in plant innate immunity was flagged (de Marchi *et al.*, 2016).

The spectrum of N-termini possibly recognized by plant N-end rule E3 ligases including PRT1 is not sufficiently explored. Only Phe-starting test substrates were found to be stabilized in a *prt1* mutant whereas initiation by Arg and Leu still caused degradation (Potuschak *et al.*, 1998; Stary *et al.*, 2003; Garzón *et al.*, 2007). In the light of substrate identification, it is cardinal to determine PRT1 mechanistics in more detail because several posttranslationally processed proteins bearing Phe, Trp and Tyr at the neo-N-termini were found (Tsiatsiani *et al.*, 2013; Venne *et al.*, 2015) and hence represent putative PRT1 targets altogether. Elucidating the substrate specificity of PRT1 will be an important step forward towards substrate identification and association of PRT1 and the N-end rule with a biological context.

We established a technique that allows real time measurements of ubiquitination using fluorescence scanning of SDS-PAGE gels and fluorescence polarization. We propose its use as a generic tool for mechanistic and enzymological characterization of E3 ligases as master components of the UPS directing substrate specificity. With a series of artificial test substrates comprising various *bona fide* destabilizing N-end rule N-termini, substrate specificity was analyzed and revealed PRT1 preference for Phe as a representative of the bulky hydrophobic class of amino acids. The methods commonly used to assay *in vitro* ubiquitination are based on end-time methods where the reaction is stopped at a given time point and analyzed by SDS-PAGE followed by immunostaining with anti-Ub versus anti-target specific antibodies. This detection via western blot often gives rise to the characteristic hallmark of polyubiquitinated proteins, a “ubiquitination smear” or a more distinct “laddering” of the posttranslationally Ub-modified target proteins. All the information of what occurred during the time of reaction is unknown unless the assay is run at several different time points which drastically increases both experimental time and reagent consumption. Besides the most common methods used for ubiquitination assessment that involve immunodetection with anti-Ub and anti-target antibodies, there are few other approaches making use of different reagents. Comparable methods, their advantages and disadvantages are listed in **Supporting Information Table S1**. The novelty offered by the present study is the development of a fluorescence-based assay that allows real-time measurement of Ub incorporation in bulky solution eliminating shortcomings of the existing methods and thus a more real mechanistic investigation. Our method monitors the ubiquitination process live, in real time, using fluorescently labeled substrate proteins and fluorescence-based detection assays, namely fluorescence polarization (FP). In addition, the protocol was coupled to fast and convenient scanning fluorescence in-gel detection. This type of assay can be easily adapted for high-throughput measurements of ubiquitination activity and probably also similar protein modification processes involving changes in substrate molecule properties over time *in vitro*. Rather than merely analyzing enzyme–substrate or protein–protein interactions, the here described method for the first time employs FP measurements for the characterization of enzyme activity and parameters affecting the performance of the ubiquitination reaction (Xia *et al.*, 2008; Kumar *et al.*, 2011; Smith *et al.*, 2013).

Here, we report a novel advanced approach to molecularly characterize E3 ligases, to measure and track polyubiquitination live and in a time-resolved manner. It has the potential to give rise to profound implications on our understanding of the interactions of E3 ligases with substrates and cofactors (non-substrates) and can impact ubiquitination research in general as our work suggests to be transferable to other E3 ligases and enzyme-substrate pairs. The method relies on rapid, easy and cheap protocols which are currently lacking for in-depth biochemical analysis of E3 ligases and is at the same time non-radioactive, sterically not interfering, and works with entire proteins in form of directly labeled substrates.

So far, only three reports mention work on PRT1 at all, i.e. the two first brief descriptions (Potuschak *et al.*, 1998; Stary *et al.*, 2003) and one highlighting the role of the N-end rule pathway, in particular a novel function for PRT1, in plant immunity (de Marchi *et al.*, 2016). However, the community lacks proofs demonstrating that PRT1 and other E3 candidates are indeed involved in substrate protein ubiquitination. To date, ubiquitination activities of E3 ligase candidates from the plant N-end rule pathway were only speculated.

## MATERIALS AND METHODS

### Cloning and expression of recombinant proteins

#### Artificial N-end rule substrates

*Escherichia coli* flavodoxin (Flv, uniprot ID J7QH18) coding sequence was cloned directly from *E. coli* DNA BL21(DE3) and flanked by an N-terminal triple hemagglutinin (HAT) epitope sequence using the primers Flv_rvs (5‘-TTATTTGAGTAAATTAATCCACGATCC-3‘) and Flv_eK_HAT(oh)_fwd (5‘-CTGGTGCTGCAGATATCACTCTTATCAGCGG-3‘). The X-eK sequences comprising codons for various N-terminal amino acids exposed after TEV cleavage of the expressed X-eK-Flv fusion protein were cloned from an eK:HAT template using the primers eK(X)_TEV(oh)_fwd (5’-GAGAATCTTTATTTTCAGxxx CACGGATCTGGAGCTTG-3’ with xxx=GTT (for Phe), GGG (for Gly), GAG (for Arg), and GTT (for Leu)) and eK_HAT_flav(oh)_rvs (5’-CCGCTGATAAGAGTGATATCTGCAGCACCAG-3’). This sequence contains a TEV protease recognition sequence (ENLYFQ|X with X being the neo-N-terminal after cleavage, i.e. TEV P1' residue) at the N-terminal of the expressed X-eK-Flv fusion protein. In order to attach Gateway attB sites and fuse the PCR products, a PCR was performed using Flv_attB2(oh)_rvs (5’-GGGACCACTTTGTACAAGAAAGCTGGGTA TCATTATTTGAG-TAAATTAATCCACGATCC-3’) and adapter_tev_fwd (5’-GGGGACAAGTTTG TACAAAAAA-GCAGGCAGGCTTAGAAAACCTGTAT TTTCAGGGAATG-3’). All primer sequences are listed in **Supporting Information Table S2**. An LR reaction into pVP16 (Thao *et al.*, 2004) (kind gift from Russell L. Wrobel, University of Wisconsin-Madison) lead to the final construct that consists of an N-terminal 8xHis:MBP double affinity tag. The expression vector pVP16::8xHis:MBP:tev:eK:3xHA:Flv was transformed into *E. coli* BL21(DE3) and the fusion protein was expressed by 0.2 mM IPTG induction in LB medium for 16 h at 26°C. Cells were harvested via centrifugation (3,500 g, 4°C, 20 min), resuspended in Ni-buffer (50 mM sodium phosphate pH 8.0, 300 mM NaCl), treated with 1mg/mL lysozyme (Sigma) in the presence of PMSF (Santa Cruz Biotechnology, sc-3597) added to a final concentration of 1 mM followed by sonication (4 min 40%, 6 min 60% intensity). The lysate was centrifuged (12,500 g, 30 min), the supernatant loaded onto a Ni-NTA agarose column (Qiagen) equilibrated with Ni-buffer, followed by Ni-buffer washing, then the protein was eluted with Ni-buffer containing 200 mM imidaziole (Merck) and loaded onto amylose resin (NEB). After washing with amylose-buffer (25 mM sodium phosphate pH 7.8, 150 mM NaCl), the protein was eluted with amylose-buffer containing 10 mM maltose. For TEV digest, the fusion protein was incubated overnight at 4°C with 0.27 µg/µL self-made TEV protease, expressed from pRK793 (Addgene, plasmid 8827), in 50 mM phosphate pH 8.0, 0.5 mM EDTA, 1 mM DTT and loaded onto a Ni-agarose column (Qiagen) equilibrated with Ni-buffer. The flow-through containing the tag-free X-eK-Flv substrate was concentrated with an Amicon Ultra-15 (Merck Millipore).

#### PRT1 cloning, expression and purification

The coding sequence of *Arabidopsis* PRT1 was cloned according to gene annotations at TAIR (www.arabidopsis.org) from cDNA. The Sequence was flanked by an N-terminal TEV recognition sequence for facilitated downstream purification using the primers ss_prt1_tev (5’-GCTTAGAGAATCTTTATTTTCAGGGGATGGCCGAAACTATGAAAGATATTAC-3’) and as_prt1_gw (5’-GGGTATCATTCTGTGCTTGATGACTCATTAG-3’). A second PCR using the primers adapter (5’-GGGGACAAGTTTGTACAAAAAAGCAGGCTTAGAGAATCTTTATTTTCAG GGG-3’) and prt1_pos2_as (5’-GGGGACCACTTTGTACAAGAAAGCTGGGTATCATTCTGTGCTT GATGA-3’) was performed to amplify the construct to use it in a BP reaction for cloning into pDONR201 (Invitrogen) followed by an LR reaction into the vector pVP16 (Thao *et al.*, 2004). Recombination into this Gateway destination vector containing a 8xHis:MBP coding sequence 5’ of the Gateway cassette leads to an N-terminal 8xHis:MBP double affinity tag. The 8xHis:MBP:PRT1 isolation, cleavage and purification was done as described above for the X-eK-Flv but the Ni-buffer contained 10% glycerol and 0.1% Tween 20.

### Chemical labeling

10 µM of purified X-eK-Flv was incubated for 1 h at room temperature with 100 µM of the synthesized thiol reactive fluorogenic labeling dye in 20 mM Tris-Cl pH 8.3, 1 mM EDTA and 1 mM tris(2-carboxy-ethyl)phosphine (TCEP, Thermo Scientific). The reaction was stopped with 1 mM cysteine hydrochloride, the unreactive dye removed using 10 kDa cut-off Amicon filters (Merck Millipore) by three successive washing steps, and the labeling efficiency evaluated by fluorescence intensity of the labeled dye (Tecan M1000) and total protein concentration using infra-red spectroscopy (Direct Detect, Merck Millipore).

### Chemical synthesis

The detailed synthesis protocols of the labeling probe NBD-NH-PEG_2_-NH-haloacetamide are described in **Supporting Information Methods**. In brief, the following synthesis steps were accomplished: 1) tert-butyl {2-[2-(2-aminoethoxy)ethoxy)ethyl}carbamate (NH_2_-PEG_2_-NHBoc); 2) NBD-NH-PEG_2_-NHBoc; 3) NBD-NH-PEG_2_-NH_2_ hydrochloride; 4) NBD-NH-PEG_2_-NH-iodo-acetamide; 5) NBD-NH-PEG_2_-NH-iodoacetamide; 6) NBD-NH-PEG_2_-NH-chloroacetamide.

### *tert*-butyl {2-[2-(2-aminoethoxy)ethoxy)ethyl}carbamate (NH_2_-PEG_2_-NHBoc)

To a solution of 2,2'-(ethylenedioxy)-bis(ethylamine) (50.00 mL, 33.83 mmol; 495.6 %) in dry dioxane (190 mL), di-*tert*-butyl dicarbonate (14.90 g, 68.27 mmol, 100 %) in dry dioxane (60 mL) was added slowly and the resulting mixture was stirred at 25 °C for 12 h. The reaction mixture was filtered, the solvent was removed under reduced pressure and the remaining residue was dissolved in distilled water (300 mL). The aqueous phase was extracted with dichloromethane (3 x 250 mL). Finally, the combined organic phases were dried (Na_2_SO_4_) and the solvent was removed under reduced pressure to yield *tert*-butyl {2-[2-(2-aminoethoxy)ethoxy)ethyl}carbamate (NH_2_-PEG_2_-NHBoc) as light yellow oil (16.09 g, 64.8 mmol, 94.9 %). ^1^H NMR (400 MHz; CDCl_3_) δ: 1.42 (br. s., 2H), 1.42 − 1.46 (m, 9H), 2.87 − 2.90 (m, 2H), 3.32 (m, 2H), 3.52 (m,, 2H), 3.55 (m,, 2H), 3.61 – 3.64 (m, 4H), 5.13 (br. s., 1H) ppm; ^13^C NMR (100 MHz, CDCl_3_) δ: 28.4, 40.3, 41.8, 67.1, 70.2, 73.5, 79.2, 156.0 ppm; ESI-MS m/z: 248.7 [M + H]^+^, 497.4 [2M + Na^+^]^+^; HRMS (ESI) calculated for C_11_H_25_N_2_O_4_ 249.1809, found 249.1809.

### NBD-NH-PEG_2_-NHB_oc_

To a suspension of *tert*-butyl {2-[2-(2-aminoethoxy)ethoxy)ethyl}carbamate (1.50 g, 6.04 mmol, 100 %) and sodium bicarbonate (1.01 g, 12.08 mmol; 200 %) in acetonitrile (30 mL), 4-chloro-7-nitrobenzofurazan (NBD) (1.80 g, 9.06 mmol, 150 %) in acetonitrile (30 mL) was added slowly over a period of 2 h and the resulting mixture was stirred at 25 °C for 12 h. The reaction mixture was filtered, the solvent was removed under reduced pressure, and the remaining residue was subjected to chromatography (silica gel, methanol / ethyl acetate, 5 : 95) to yield NBD-NH-PEG_2_-NHB_oc_ as a brown solid (1.89 g, 4.58 mmol, 75.9 %). M.p.: 85 – 86 °C; *R* _F_ = 0.56 (methanol / ethyl acetate, 5 : 95); ^1^H NMR (400 MHz; CDCl_3_) δ [ppm]: 1.42 – 1.45 (m, 9H), 3.31 – 3.37 (m, 2H), 3.54 – 3.56 (m, 2H), 3.58 – 3.60 (m, 2H), 3.61 – 3.71 (m, 4 H), 3.87 (m, 2H), 5.02 (m, 1H), 6.20 (d, *J* = 8.6 Hz, 1H), 6.88 (m, 1H), 8.49 (d, *J* = 8.6 Hz, 1H); ^13^C NMR (100 MHz; CDCl_3_) δ [ppm]: 28.4, 43.6, 68.1, 70.2, 70.2, 70.4, 70.5, 77.2, 98.7, 136.3, 143.9, 144.0, 144.0, 144.3, 155.9; ESI-MS m/z: 410.5 [M − H]^+^, 434.2 [M + Na]^+^, 845.4 [2M + Na]^+^; HRMS (ESI) calculated for C_17_H_25_N_5_O_7_Na 434.1646, found 434.1647.

### NBD-NH-PEG_2_-NH_2_ hydrochloride

To a solution of NBD-NH-PEG_2_-NHBoc (2.08 g, 5.06 mmol, 100 %) in dry methanol (20 mL), trimethylsilyl chloride (2.70 mL, 21.27 mmol, 500 %) was added *via* syringe and the resulting mixture was stirred at 25 °C for 12 h. The solvent was removed under reduced pressure. The remaining residue was suspended in diethyl ether (15 mL), filtered and the solid was washed with several portions of diethyl ether, and the remaining solid was dried under reduced pressure to yield NBD-NH-PEG_2_-NH_2_ hydrochloride as a brown solid (1.56 g, 5.01 mmol, 98.9 %). The crude product was used without further purification. M.p.: 192 – 193 °C; ^1^H NMR (400 MHz; CD_3_OD) δ [ppm]: 3.09 – 3.11 (m, 2H), 3.64 – 3.76 (m, 8H), 3.87 – 3.90 (m, 2 H), 6.19 (d, *J* = 8.4 Hz, 1H), 8.45 (d, *J* = 8.7 Hz, 1H); ^13^C NMR (100 MHz; CD_3_OD) δ [ppm]: 41.5, 41.7, 70.1, 70.3, 70.8, 73.2, 98.8, 123.0, 136.5, 144.1, 144.4, 144.8; ESI-MS m/z: 310.5 [M − 2H]^+^, 312.3 [M]^+^; HRMS (ESI) calculated for C_12_H_18_N_5_O_5_ 312.1303, found 312.1303.

### NBD-NH-PEG_2_-NH-iodoacetamide

To a solution of NBD-NH-PEG_2_-NH_2_ hydrochloride (202.3 mg, 0.65 mmol; 100 %) and *N*,*N* '-diisopropylethylamine (134.3 µL, 0.77 mmol, 120 %) in dry acetonitril (4.0 mL), iodoacetic anhydride (401.0 mg, 1.13 mmol; 174 %) was added slowly and the resulting mixture was stirred at 25 °C for 12 h. The solvent was removed under reduced pressure and the remaining residue was subjected to chromatography (silica gel, methanol / ethyl acetate, 10 : 90) to yield NBD-NH-PEG_2_-NH-iodoacetamide as a brown solid (151.1 mg, 0.32 mmol, 48.5 %). *R*_F_ = 0.45 (methanol / ethyl acetate, 10 : 90); ^1^H NMR (400 MHz; CDCl_3_) δ [ppm]: 3.50 – 3.54 (m, 2H), 3.62 – 3.65 (m, 2H), 3.69 – 3.71 (m, 8H), 3.73 – 3.76 (m, 2H), 6.21 (d, *J* = 8.7 Hz, 1H), 6.55 (br. s., 1H), 6.95 (br. s., 1H), 8.48 (d, J = 8.6 Hz, 1H); ^13^C NMR (100 MHz; CDCl_3_) δ [ppm]: 0.56, 40.1, 43.6, 68.1, 69.4, 70.3, 70.5, 136.4, 143.9, 144.3, 167.1; ESI-MS m/z: 478.3 [M – H]^+^, 502.1 [M + Na]^+^ + 981.3 [2M + Na]^+^; HRMS (ESI (negative modus)) calculated for C_14_H_17_N_5_O_6_I 478.0229, found 478.0222.

### NBD-NH-PEG_2_-NH-chloroacetamide

To a solution of NBD-NH-PEG_2_-NH_2_ hydrochloride (202.5 mg, 0.65 mmol; 100 %) and *N*,*N* '-diisopropylethylamine (134.3 µL, 0.77 mmol, 120 %) in dry acetonitril (4.0 mL), chloroacetic anhydride (221.7 mg, 1.30 mmol; 200 %) was added slowly and the resulting mixture was stirred at 25 °C for 12 h. The solvent was removed under reduced pressure and the remaining residue was subjected to chromatography (silica gel, methanol / ethyl acetate, 10 : 90) to yield NBD-NH-PEG_2_-NH-chloroacetamid as a brown solid (150.5 mg, 0.39 mmol, 59.7 %). *R*_F_ = 0.46 (methanol / ethyl acetate, 10 : 90); ^1^H NMR (400 MHz; CDCl_3_) δ [ppm]: 3.54 – 3.58 (m, 2H), 3.64 – 3.75 (m, 8H), 3.87 – 3.90 (m, 2H), 4.06 (m, 2H), 6.20 (d, *J* = 8.7 Hz, 1H), 6.90 (br. s., 1H), 6.98 (br. s., 1H), 8.48 (d, *J* = 8.6 Hz, 1H); ^13^C NMR (100 MHz; CDCl_3_) δ [ppm]: 30.51, 42.7, 43.6, 68.1, 69.5, 70.3, 70.5, 136.3, 143.9, 144.3, 166.0; ESI-MS m/z: 386.1 [M – H]^+^, 410.1 [M + Na]^+^; HRMS (ESI (negative modus)) calculated for C_14_H_17_N_5_O_6_Cl 386.0873, found 386.0863.

### Ubiquitination assay and in-gel fluorescence detection

3.4 µM (total protein concentration, both label and unlabeled) of the X-eK-Flv fluorescently labeled substrate (X-eK-Flv-NBD) were solved in 25 mM Tris-Cl pH 7.4, 50 mM KCl, 5 mM MgCl_2_, 0.7 mM DTT containing 16 µM Ub from bovine erythrocytes (Sigma-Aldrich, U6253). For ubiquitination, 2 mM of ATP (New England Biolabs), 40 nM of E1^15^, 0.31 µM of E2 (UBC8)^15^, and 5 nM of E3 (8xHis:MBP-tagged or untagged PRT1) were added to the previous mix in a final volume of 30 µL and incubated at 30°C for 1 h. The reaction was stopped by adding 5X reductive SDS-PAGE loading buffer and incubating for 10 min at 96 °C followed by SDS-PAGE. The gels were scanned using fluorescence detection on a Typhoon FLA 9500 biomolecular imager (GE Healthcare) with a blue excitation laser (473 nm) LD and an LBP emission filter (510LP), then blotted onto a cellulose membrane and detected with either mouse monoclonal anti-Ub antibody (Ub (P4D1), sc-8017, Santa Cruz Biotechnology, 1:5,000 dilution in blocking solution [150 mM NaCl, 10 mM Tris-Cl pH 8, 3% skim milk powder, 0.1% Tween 20]) or mouse monoclonal anti-HA epitope tag antibody (HA.11, clone 16B12: MMS-101R, Covance; 1:1,000 to 1:5,000, in blocking solution) and goat anti-mouse IgG-HRP (1858415, Pierce; 1:2,500 to 1:5,000 dilution in blocking solution). The acquired images of the gels (prior blotting) were analyzed using the Gel Analyser densitometric soft (Gel.Analyser.com). Thus, one may use the same gel for both in-gel fluorescence detection followed by blotting and immunodetection.

The same gels that were detected via fluorescence scanning were blotted and detected with ECL without further processing such as stripping. Thus, fluorescent detection can be combined with ECL in one simple workflow. For evaluation of pH dependence, 50 mM Tris-Cl was used as a buffering agent at pH 6.75, 7.0, 7.5, 8.0, 8.5 and 9.0.

### Real-time ubiquitination assay using fluorescence polarization

For fluorescence polarization (FP), the reaction mixture (24 µL) containing all the components except the ATP was incubated in a 384 well microplate (Corning, Cat. No. 3712 or 3764) at 30°C in a M1000 infinite plate reader (Tecan) until the temperature was stable (typically 4-5 min) and the reaction triggered by adding 6 uL of 10 mM ATP preheated to 30°C. FP was monitored every 2 min at 562 nm while the excitation wavelength was set to 470 nm. The M1000 fluorescence polarization module was calibrated using 10 nM fluorescein in 10 mM NaOH at P = 20 mP.

### Structure modeling of the artificial substrate

The amino acid sequence of the artificial F-eK-Flv substrate was submitted to the Protein Homology/Analogy Recognition Engine V 2.0 (Kelley *et al.*, 2015) (Phyre^2^, Structural Bioinformatics Group, Imperial College, London) in both normal and intensive modes. The bests selected templates were found to be PBD ID: 3EDC for the eK region and 2M6R for the Flv part) and the model was visualized using ViewerLite (Accelrys Inc.).

## RESULTS

### PRT1 is an E3 ubiquitin ligase and prefers bulky N-termini

For the analysis of PRT1 E3 ligase function, i.e. recognition of N-end rule substrates, we used recombinant PRT1 together with generic substrate reagents with unprecedented detection features combining chemically synthesized fluorophores and recombinant ubiquitination acceptors which were used as live protein modification detectors. To describe N-terminal amino acid specificity of PRT1, the N-terminally variable protein parts of the reporters were engineered as N-terminal His8:MBP fusions comprising a recognition sequence of tobacco etch virus (TEV) protease at the junction to the subsequent generic substrate protein moiety (**Fig. 2a, Supporting Information Fig.S 1a**). Cleavage by TEV gave rise to small C-terminal fragments of the His8:MBP-substrate fusions of which the neo-N-terminal, i.e. the P1' residue of the TEV cleavage site, can be altered to all proteinogenic amino acids except proline (Kapust *et al.*, 2002; Phan *et al.*, 2002; Naumann *et al.*, 2016). For a novel fluorescence-based approach, we covalently coupled a synthetic fluorescent probe (**Fig. 2b**) to the artificial substrate protein. The resulting reagent served as fluorescent protein Ub acceptor in N-end rule ubiquitination assays. The architecture of the reagent is as follows: after the cleavable His8:MBP tag, eK, a part of *E. coli* lacZ (Bachmair *et al.*, 1986) followed by a triple hemagglutinin epitope tag (3HA) for immunodetection and an *E. coli* flavodoxin (Flv) were combined. Flv was chosen as a highly soluble and stable protein and includes flavin mononucleotide as a cofactor. Its semiquinone is fluoresent but not stable enough to be used as fluorophore for detection in its plain form. Therefore, we decided to additionally label the Flv protein. The junctions between His8:MBP and eK encode for the N-termini glycin (Gly, G), phenylalanin (Phe, F), arginine (Arg, R), and leucin (Leu, L) that get N-terminally exposed after TEV cleavage. The G/F/L/R-eK-Flv constructs contain a single cysteine (Cys101 of Flv) that allowed the labeling of the purified recombinant fusion protein with a novel thiol-reactive probe that comprises an iodoacetamide-polyethylene glycol (PEG) linker and the fluorogenic subunit of 4-nitro-2,1,3-benzoxadiazole (NBD; **Fig. 2b**). We chose the latter due to its small size compared to other labeling reagents such as large fluorescein moieties and because it can be detected very specifically by both UV absorption and UV fluorescence with low background interferences.

**Figure 2.**
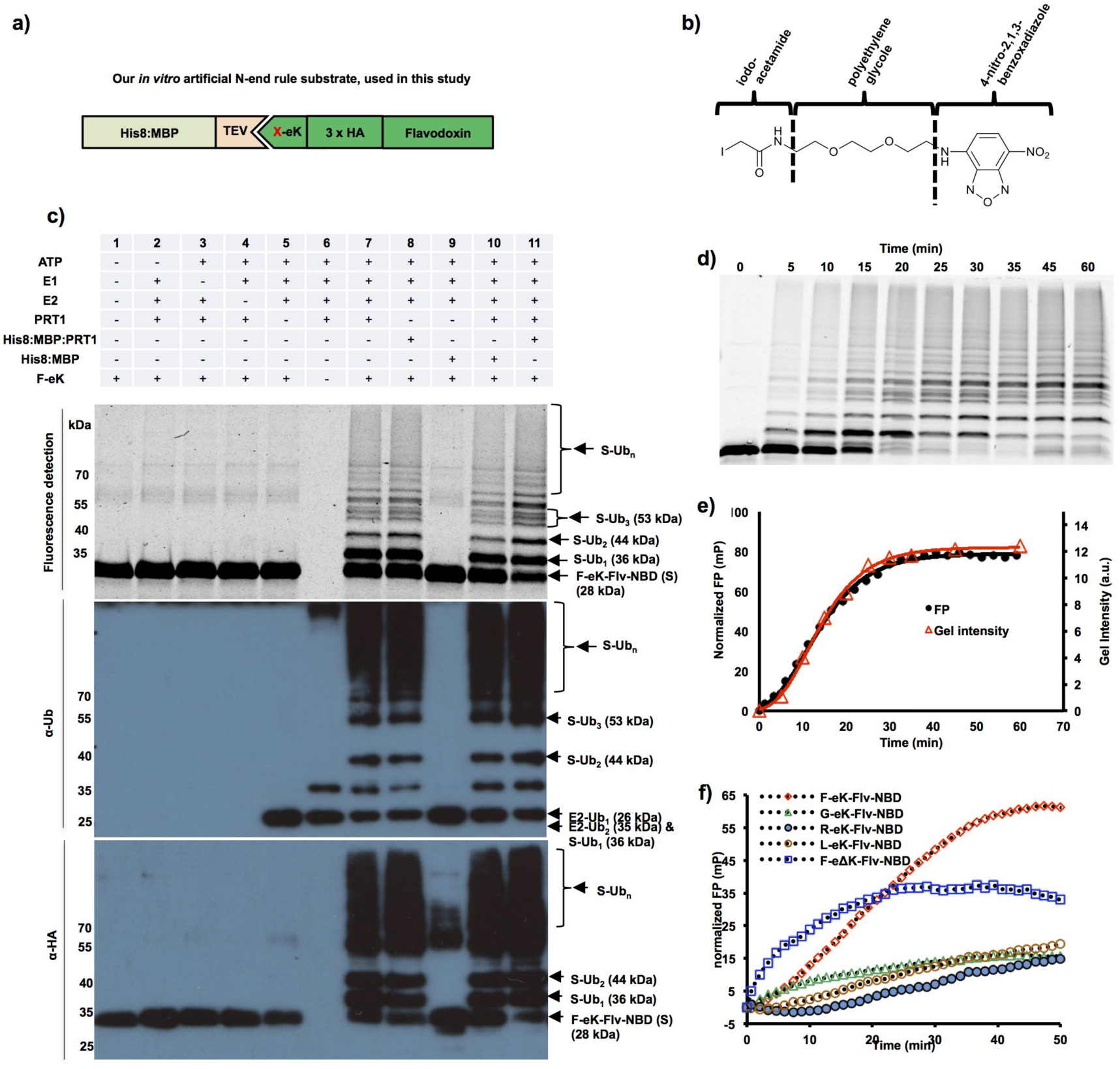
Fluorescent protein conjugates for monitoring *in vitro* substrate ubiquitination in real time. **a)** Design of recombinant fusion proteins used as N-end rule substrates. After TEV cleavage and removal of the His8:MBP affinity tag, the artificial substrate based on *E. coli* flavodoxin (Flv) is initiated with a neo-N-terminal, here Phe (F), Gly (G), Leu (L) or Arg (R). **b)** Skeletal formula of the synthesized thiol-reactive fluorescent compound. The substrate was covalently tagged with the reagent composed of iodoacetamide, polyethylene glycol (PEG) linker and 4-nitro-2,1,3-benzoxadiazole (NBD). The reactive iodine-containing group on the left couples to the thiol group of internal Cys residues of Flv. NBD serves as a fluorophore with excitation at 470 nm and emission at 520 nm. **c)** Detection via fluorescence and immunoblotting of the F-eK-Flv-NBD after *in vitro* ubiquitination. The labeled protein and its ubiquitinated variants were detected via fluorescence scanning directly from the SDS-PAGE gel followed by western blotting and immunodetection with anti-HA and anti-Ub antibodies. Lane 6 shows ubiquitinated E2 like in all lanes and autoubiquitination of PRT1 as very high molecular weight ‘smear’. Cleaved PRT1 as well as His8:MBP-tagged PRT1 were used together with His:UBA1 (E1) and His:UBC8 (E2) (Stegmann *et al.*, 2012). **d and e)** Kinetic profiles of PRT1-mediated ubiquitination. F-eK-Flv-NBD ubiquitination was monitored by FP and in-gel fluorescence scanning. The S-shaped kinetic curve is observed in both in-gel fluorescence scanning detection and fluorescence polarization. **f)** N-terminal specificity evaluated by real-time ubiquitination detection. Fluorescently labelled R-eK-Flv, L-eK-Flv, G-eK-Flv, F-eΔK-Flv and F-eK-Flv were comparatively evaluated for their degree of ubiquitination by PRT1.

In an *in vitro* ubiquitination assay, we used recombinant UBC8 as a promiscuous E2 conjugating enzyme and UBA1 as E1 activating enzyme (Stegmann *et al.*, 2012) and show here for the first time E3 ligase activity of PRT1 depending on E1, E2 and ATP (**Fig. 2c**). PRT1 discriminated a substrate by its N-terminal, aiding the transfer of Ub to the substrate and leading to polyubiquitination. After immunostaining with anti-Ub antibodies, usually, a typical smear of higher molecular weight compared to the target protein's size is observed or after probing with target-specific antibodies, a more or less distinct laddering, also of high molecular weight, becomes evident. These are the common signs for polyubiquitination and a clear laddering was also visualized by fluorescent scanning in our novel approach. We identified distinct subspecies via in-gel detection (**Fig. 2c**). A classical end-time point assay where the reaction was stopped at different reaction time points followed by SDS-PAGE and in-gel fluorescence detection revealed the kinetics of PRT1 activity using F-eK-Flv as substrate (**Fig. 2d**).

However, a real-time monitoring of the kinetic profile of the enzymatic reaction is only possible via FP in live detection measurements. The kinetic profile is best-fitted with an S-shaped curve and a growth curve model of logistic type (Richards’ equation) rather than exponentially as expected for simple kinetics (**Fig. 2e**).

It was previously suggested that PRT1 binds to N-degrons carrying bulky destabilizing residues (Stary *et al.*, 2003) but biochemical evidence for that was still lacking. By changing the N-terminal residue of the X-eK-Flv-NBD substrate, it was possible to reveal that PRT1 indeed discriminates the substrates according to the N-terminal residue, as expected (**Fig.2f, Supporting Information Fig. S1b,c**). While the substrates carrying G-, R-, L-initiated N-termini showed poor ubiquitination, F-eK-Flv-NBD was heavily ubiquitinated. While the eK-based substrate showed the kinetic curve discussed above, the control F-eΔK-Flv substrate with mutated lysines (expected site of ubiquitination, Lys15 and Lys17, both replaced by Arg) presented a faster initial rate of ubiquitination but levels of only half of the final FP value (**Fig. 2f**). This is in good agreement with the in-gel fluorescence detection where lower degrees of ubiquitination of F-eΔK-Flv, reduced mono- and di-ubiquitination - but still clear polyubiquitination - were observed (**Supporting Information Fig. S1c**).

Another remarkable observation of the ubiquitination pattern in the in-gel fluorescence image (using three different independent substrate protein purifications of F-eK-Flv-NBD) was that the tri-ubiquitinated form presents three distinct subspecies which eventually lead to a multitude of other species at higher level (**Supporting Information Fig. S1b**). There was only one species of tri-ubiquitinated F-eΔK-Flv-NBD generated, where two ubiquitination acceptors sites within eK (Lys15 and Lys17) were replaced by Arg (**Supporting Information Fig. S1b**).

### Fluorescently labeled substrate proteins unravel mechanism of PRT1-mediated ubiquitination

The combination of the proposed two fluorescence-based methods allowed fast and efficient *in vitro* investigation of the ubiquitination process via the E3 ligase PRT1 and the optimization of the reaction conditions. As a first approach utilizing the real-time assay in the context of substrate ubiquitination, we studied the role of changes in pH on the ubiquitination process mediated by PRT1. A classical end-time approach revealed the reaction optimum to be clearly above pH 7 but below pH 9 as indicated by the occurrence of polyubiquitinated species of the fluorescent substrate probe F-eK-Flv-NBD (**Fig. 3a**). However, using our real-time FP protocol, we additionally acquired the kinetic profile of the PRT1-mediated ubiquitination process (**Fig. 3b**) and the maximum reached polarization values of this reaction (**Fig. 3c**). These correlated with the amount of polyubiqutinated species detected in the SDS-PAGE gel-based end-time experiment (**Fig. 3a**) and the highest initial rate (**Fig. 3c**) whereas the latter appears to be different from the reaction optimum according to the detected max. FP. We also had previously observed, that F-eΔK-Flv ubiquitination presented a faster initial rate but only half of the final FP (**Fig. 2f**) and lower degrees of final ubiquitination (**Supporting Information Fig. S1c**). Both bell-shaped forms of the pH dependence for the highest initial reaction rate (pH 8.0) and the maximum substrate polyubiquitination rate (pH 7.5) indicated two competing processes that generate a local maximum (**Fig. 3c**).

**Figure 3.**
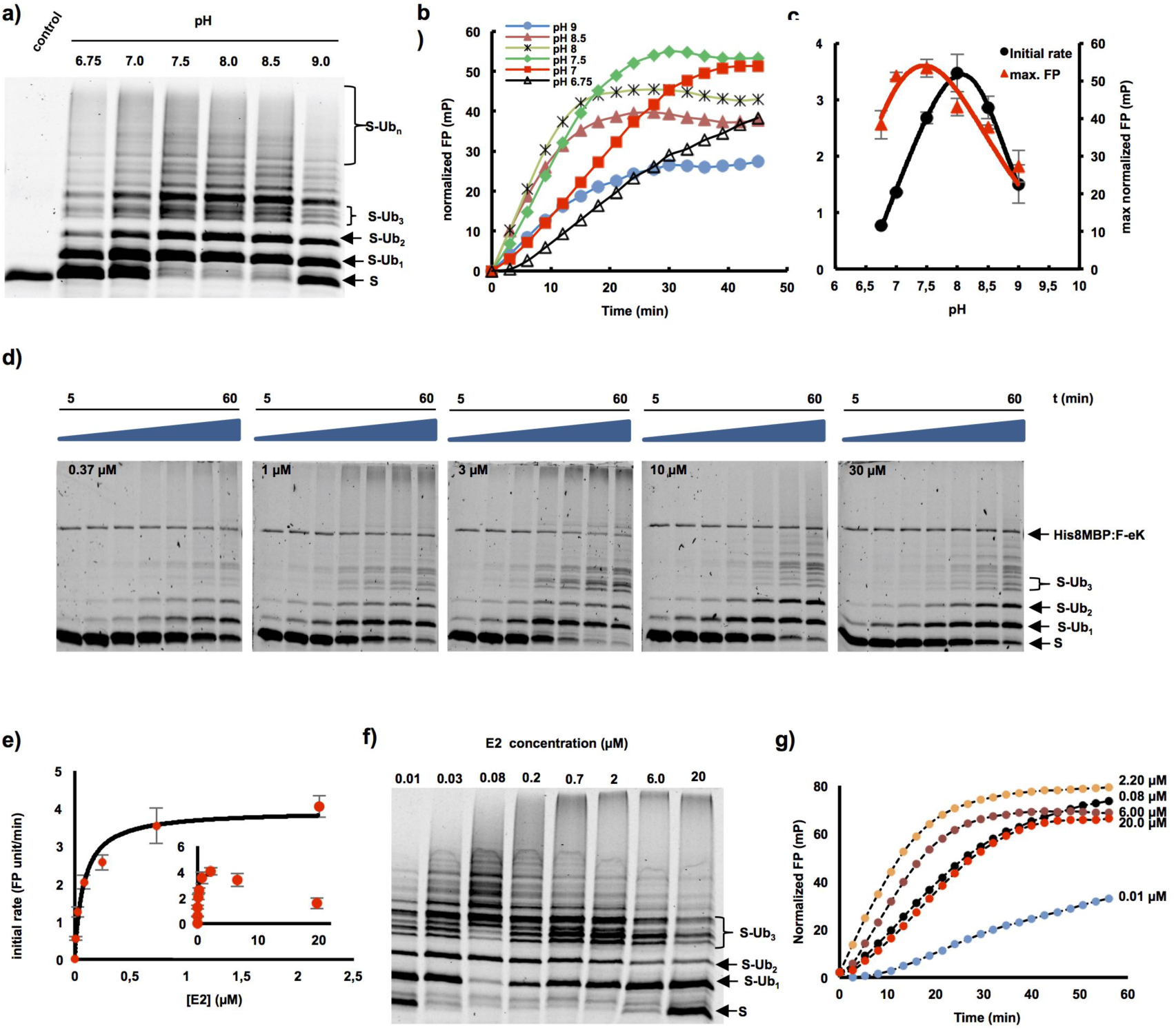
Applications of fluorescent protein conjugates for monitoring pH dependent ubiquitination and enzymatic parameters of PRT1 E3 ligase. **a-c)** pH dependent ubiquitination of the F-eK-Flv substrate. **a)** In-gel detection of F-eK-Flv ubiquitinated species after 1 h reaction at several pH values demonstrating different patterns of polyubiquitination preferences depending on the pH. **b)** Kinetic profiles, **c)** initial rates and maximum end-time FP values forming a bell-shaped distribution depending on the pH. **d-g)** PRT1-mediated ubiquitination of F-ek-Flv dependent on the concentration of E2-conjugating enzyme (UBC8). **d)** Time dependence of ubiquitination at several E2 concentrations for the first 60 min at 5 nM PRT1, time scale: 5-60 min. **e)** Michaelis-Menten curve plotted using the initial rate from FP data suggest an E2-driven inhibition effect. **f)** The qualitative evaluation of ubiquitination was done using in-gel scanning fluorescence and **g)** kinetic profiles were obtained using FP measurements, similar conditions as in d) but with ten times higher concentration of PRT1, i.e. 50 nM.

A strong decrease of the ubiquitination rate mediated by PRT1 was observed at higher concentrations of the E2-conjugating enzyme UBC8 (>2 µM) both via in-gel fluorescence (**Fig. 3d**) and FP (**Fig. 3e-g**). Based on the FP measurements using up to 2 µM of UBC8, the KM of substrate ubiquitination by PRT1 at different E2 concentrations was found to be in the submicromolar range, 0.08±0.01 µM, indicating a very tight binding of the E2 to PRT1 compared to other RING E3 ligases (Ye & Rape, 2009) (**Fig. 3e**). Moreover, the distribution pattern of the ubiquitinated substrate species at the end of the reaction (**Fig. 3f**) and the kinetic profiles of ubiquitination (**Fig. 3g**) are different, depending on the used E2 concentration.

## DISCUSSION

The N-end rule pathway is an emerging vibrant area of research in plant sciences and agriculture (Gibbs *et al.*, 2011; Licausi *et al.*, 2011; Gibbs *et al.*, 2014b; Weits *et al.*, 2014; de Marchi *et al.*, 2016; Mendiondo *et al.*, 2016) and reviewed in (Gibbs *et al.*, 2014a; Gibbs, 2015; Gibbs *et al.*, 2016). Taking the first *bona fide* plant N-end rule E3 Ub ligase PRT1 as the model, we describe a novel tool to molecularly characterize polyubiquitination live, in real-time, and use it to gain the first mechanistic insights in PRT1 substrate preference, activation and functional pairing with an E2-conjugating enzyme. To date, activity and function of enzymatic N-end rule pathway components was only speculated and the field was lacking investigations on molecular level. Here, we showed the first molecular evidence for ubiquitination activity of an E3 ligase candidate from the entire plant N-end rule pathway.

Here, we demonstrated PRT1 E3 Ub ligase activity and substrate preference by using recombinant PRT1 together with artificial protein substrates in an *in vitro* fluorescence-based life ubiquitination assay. We found that first of all, the reporter construct based on bacterial Flv chemically coupled to NBD (**Fig. 2b**) works as ubiquitination acceptor. Second, this reaction reflects substrate specificity and cannot be considered an *in vitro* artifact, since N-terminal amino acids other than Phe rendered the substrate a weaker target for PRT1 (**Fig. 2f, Supporting Information Fig. S1b,c**). Third, our test system allowed to describe E3 ligase function and target specificity by using variants of labeled substrates.

Similar experiments are usually evaluated based on immunochemical and colorimetric detection, incorporation of radioisotopes such as ^125^I or ^32^P, or fluorescently labeled native or recombinant Ub (Ronchi & Haas, 2012; Melvin *et al.*, 2013; Lu *et al.*, 2015a; Lu *et al.*, 2015b) (**Supporting Information Tab. S1**). However, problems of steric hindrance by modifying Ub and difficulties to discriminate between auto- and substrate ubiquitination if using labeled Ub may occur. Also artificial experimental setups such as single-molecule approaches or extreme buffer conditions might not represent or support formation of the required complex ubiquitination machinery (**Supporting Information Table 1**). Our assay allowed both direct assessment in the flow of the actual FP experiment and gel-based evaluation after completing SDS-PAGE. This renders protein transfer via western blotting plus the subsequent time-consuming steps of blocking, immuno- and chemical detection obsolete. The protocol described is rapid, non-radioactive, uses only a small fluorophore as a covalent dye, works with full substrate proteins instead of only peptides, and can be read out live in real-time. Moreover, the FP approach conveys superimposable kinetic curves with data from classical end-time point assays, but faster, with higher resolution in time and using fewer reagents. The advantage of a combination of the described two fluorescence-based approaches, that is, gel-based and FP, is the possibility to gain mechanistic insights which was not possible by applying only one of the single protocols. An example is the determination of KM and kcat of the interaction of the E3 ligase PRT1 with E2-conjugating enzymes. This included the influence of the E2 concentration on both the ubiquitinated substrate species and the kinetic profile of the ubiquitination reaction.

Using FP coupled to immunoblot analysis, we were able to confirm that PRT1 is an active E3 ligase acting in concert with E2-conjugating enzyme UBC8. In a buffer system close to physiological conditions, it could be shown that PRT1 not only monoubiquitinates N-degron containing substrates, but also mediates polyubiquitination without the aid of further cofactors. Therefore, it was ruled out that PRT1 only monoubiquitinates which was speculated previously (Stary *et al.*, 2003). Moreover, the action of a type II-N-recognin as small as PRT1 (46 kDa) is most likely sufficient for subsequent target degradation by the proteasome. Since PRT1 lacks the conserved ClpS domain that confers affinity to type II substrates in other N-recognins, the binding mechanism of PRT1 to its substrate remains an intriguing open question.

By FP-facilitated real-time monitoring of the kinetic profile of the PRT1-mediated ubiquitination, we observed the S-shaped curve of the reaction (**Fig. 2e**). One explanation for this kinetics and the presence of an initial lag phase is an increase of the affinity of PRT1 for the monoubiquitinated substrates compared to the non-ubiquitinated population. Preferences of E2s and E3s for mono- or polyubiquitinated substrates and their influence on ubiquitination velocity but also that initial ubiquitination greatly enhances the binding affinity of E3s to the substrate in subsequent reactions was shown previously (Sadowski & Sarcevic, 2010; Lu *et al.*, 2015b). The chain elongation (Ub-Ub isopeptide bond formation) can be faster than the chain initiation which might represent the rate limiting-step of the reaction, rather than an E1-E2-controlled limiting-step. Thus, the chain elongation and chain initiation steps appear to be distinct processes that have distinct molecular requisites in agreement with previous findings for other E3s (Petroski & Deshaies, 2005; Deshaies & Joazeiro, 2009). The lag phase is reduced if the rate is increased by higher concentration of PRT1 (**Fig. 2e**).

The FP-based assay revealed that the kinetic profile of the ubiquitination was dependent on the position and availability of lysines as Ub acceptor sites as suggested to be characteristic of N-degrons (Bachmair & Varshavsky, 1989). By lowering the overall number of available lysines in the F-eΔK-Flv-NBD substrate (two Lys less than in X-eK-Flv constructs with 11 Lys in total) the overall ubiquitination was detectably reduced. Differences in the kinetic curves of F-eK-Flv versus F-eΔK-Flv indicated that a reduction of the available number of Lys residues lead to a faster initial rate of ubiquitination whereas the final FP values reached only half of the levels compared to the assay applying the substrate with the full set of Lys residues (**Fig. 2f**, **Supporting Information Fig. S1c**). However, the simple gel-based end-point assay could not unravel if this was due to altered velocity of chain initiation versus chain elongation. The initiation per Lys residue was expected to be similar in F-eK- versus F-eΔK-Flv substrates but chain elongation could apparently start faster in F-eΔK-Flv. This demonstrated that the presence of E2 together with the particular substrate plays a key role in the formation of the molecular assembly facilitating the ubiquitination process. Already the intermolecular distance between the E3 ligase and the Ub acceptor lysines of the substrate as well as the amino acid residues proximal to the acceptor lysines determine the progress of the reaction and ubiquitination specificity (Sadowski & Sarcevic, 2010). Taking the slower initiation of polyubiquitination of F-eK-Flv into account, the availability of lysines at the N-terminus might interfere with the monoubiquitination of other, more distal lysines and the E3 could remain associated with substrates that are monoubiquitinated at the N-terminal.

When subjecting the F-eK-Flv-NBD substrate fusion protein to in vitro ubiquitination assays, three distinct subspecies of the tri-ubiquitinated form were detected versus only one form, if F-eΔK-Flv-NBD was used (**Supporting Information Fig. S1c**). This could be explained by a formation of various ubiquitinated isoforms of the substrate by utilizing different lysine side chains as ubiquitination acceptor sites. These could be either within the sequence of eK (e.g. Lys15 and Lys17) or within Flv (e.g. Lys100 and Lys222 which seem structurally more favored according to the structural model, **Supporting Information Fig. S1a**). This was further supported by the fact that there is only one species of tri-ubiquitinated F-eΔK-Flv-NBD, where two ubiquitination acceptor sites within eK (Lys15 and Lys17) were replaced by Arg (**Supporting Information Fig. S1b**).

When analyzing the influence of the pH on PRT1 function as E3 Ub ligase, we documented bell-shaped forms of the pH dependence for the highest initial reaction rate (pH 8.0) and determined the maximum substrate polyubiquitination rate (pH 7.5). These indicated two competing processes that generate a local maximum (**Fig. 3c**). In the light of recently discussed mechanisms of E3 ligase action (Berndsen & Wolberger, 2014) and the prediction of two RING domains in PRT1 (Stary *et al.*, 2003), higher ubiquitination rates with increased pH could be due to deprotonation of the attacking lysine side chain of the E2 active site. This would facilitate thioester cleavage between E2 and Ub and thereby mediate Ub transfer to the substrate lysines. A similar effect was observed regarding the influence of the acidic residues in close vicinity of the E2 active site, which also cause deprotonation of the lysine side chain of the incoming substrate (Plechanovova *et al.*, 2012). This possibly explains the drastic increase in the initial rate of PRT1 substrate ubiquitination in the pH 6.8 to pH 8 range (**Fig. 3c**). The competing processes leading to the decrease in ubiquitination at pH>8 could be destabilization of ionic and hydrogen bonds at alkaline pH simply interfering with protein-protein interaction or ATP hydrolysis affecting the Ub charging of the E2 by the E1. This could also explain the premature leveling of the kinetic curves in the FP measurements at pH>8 (**Fig. 3b**) while in a longer reaction timescale, the maximum FP values would be expected to be the same from pH 6.8 to pH 7.5.

The apparent catalytic rate constant (kcat) of the Ub transfer, more precisely the transfer of the first Ub molecule, i.e. the rate limiting step, was found to be 1.30±0.07 s^-1^. This suggested that on the one hand PRT1 had a high turnover number due to a highly active catalytic center and on the other hand that the E2 concentration does not only influence the rate of the Ub transfer to the substrate but also the mechanism itself. Possible causes are the two separate and potentially distinctly favored chain initiation and elongation processes mentioned above. These could result in lowering the rate of the initiation step at higher E2 concentrations since both the kinetic profile and the formation of ubiquitinated species are affected and also the attacking lysines might be structurally differently favored. This is especially suggested by the variable occurrence of the distinct pattern of triubiquitinated substrate species (**Fig. 3d,f**) as mentioned above and discussed in other systems as well (Ye & Rape, 2009).

By using fluorescently labeled substrate proteins in the two described approaches, that is, gel-based fluorescence scanning after SDS-PAGE and FP, we were able to investigate the mechanism of PRT1-mediated ubiquitination and optimize the reaction conditions. The presented work serves as a model for the demonstration of differential mechanisms of substrate recognition and tight interactor-binding in the N-end rule pathway.

PRT1 is a plant pioneer enzyme lacking homologs in the other kingdoms albeit small and easy to produce in an active form as recombinant protein rendering it an exciting candidate for further functional and structural studies of key functions of one branch of the N-end rule pathway. So far, only three research articles mention work on PRT1, i.e. the two first brief descriptions (Potuschak *et al.*, 1998; Stary *et al.*, 2003) and one recently published study highlighting the role of the N-end rule pathway - and in particular a novel function for PRT1 - in plant immunity (de Marchi *et al.*, 2016). However, to date, the community lacks proofs demonstrating that PRT1 and other E3 candidates are indeed involved in substrate protein ubiquitination.

The here described tool can be adopted by laboratories investigating N-end rule related posttranslational modifications such as deformylation, methionine excision, oxidation, deamidation, arginylation, ubiquitination and degradation. Moreover, we are convinced that it may also be extended to assays for other posttranslational modifications such as phosphorylation and to other E3 Ub ligases as long as at least one native or artificial substrate protein for the modification of interest is known. Because it makes use of chemical labeling of substrate proteins rather than labeling protein modifiers such as Ub or phosphate themselves, one common reagent can be used for various modification assays. The approach allows to measure and track posttranslational protein modification live and in a time-resolved manner and has profound implications for our understanding of the interactions of E3 ligases with substrates and non-substrates. Concerning the field of the N-end rule pathway, this might apply to other candidates of E3 Ub ligases such as PROTEOLYSIS6 (PRT6) and BIG (AT3G02260) or potential N-end rule adapter proteins like PRT7 (AT4G23860) (Tasaki *et al.*, 2005; Garzón, 2008; Talloji, 2011). These experiments will be of premier interest in the future because phenotypes of biological importance and genetically determined causalities were described and need to be substantiated on molecular level. Therefore, we see potential for a broader impact for ubiquitination research as it is conceivable that the method is transferable to other E3 ligases and enzyme-substrate pairs. In the course of our studies, we felt that rapid, easy and cheap protocols were lacking for in-depth biochemical analysis of E3 ligase kinetics, the same holds true for non-radioactive and sterically not interfering protocols and those where entire proteins and directly labeled substrates can be applied.

In terms of further applications, the kinetic approach allowed collecting data that can assist to set up high-throughput assays, e.g. for screens of inhibitors and the influence of small molecules potentially facilitating or enhancing ubiquitination. In our example, this included testing of the enzymatic parameters of E2-E3 interactions and substrate specificities for PRT1. Similar approaches have used labeling with radionuclides or fluorescent dyes coupled to Ub (Ronchi & Haas, 2012; Melvin *et al.*, 2013; Lu *et al.*, 2015a; Lu *et al.*, 2015b). The latter covalent modification of Ub with fluorescent moieties is often impractical since these groups can sterically hinder the E1-catalyzed activation and E2-dependent transthiolation reactions (Ronchi & Haas, 2012). This in turn can alter the rate-limiting step. The use of radioactive isotopes requires at least running an SDS-PAGE and gel-drying or western blotting followed by autoradiography for hours to days (**Supporting Information Tab. 1**). Besides the described *<u>in vitro</u>* methods, several protocols and tools were successfully applied *in vivo*, mainly based on translational fusions of fluorescent proteins to degrons of the Ub fusion degradation (UFD) pathway (Hamer *et al.*, 2010; Matilainen *et al.*, 2016), the N-end rule pathway (Speese *et al.*, 2003; Faden *et al.*, 2016) or both (Dantuma *et al.*, 2000). Other methods make use of Ub-binding systems to achieve various read-outs (Marblestone *et al.*, 2012; Matilainen *et al.*, 2013)(**Supporting Information Tab. 1**).

In conclusion, we describe a system for real-time measurements of ubiquitination in bulky solution with combined fluorescence scanning of SDS-PAGE gels and fluorescence polarization. This setup was used to establish an artificial substrate protein-based detection reagent that reveals important mechanistic insights of E2-PRT1-substrate interaction. We demonstrate for the first time that PRT1 is indeed involved in polyubiquitination of substrate proteins depending on its N-terminal amino acid and therefore approached PRT1 as an player of the N-end rule pathway for the first time on a molecular level.

## ACKNOWLEDGEMENTS

We thank Marco Trujillo for expression clones of His-tagged UBC8 and UBA1, discussions and constant support in ubiquitination-related issues and Angela Schaks for synthesis of the chemical probe. This work was supported by a grant for setting up the junior research group of the *ScienceCampus Halle – Plant-based Bioeconomy* to N.D., by the grant WE 1467/13-1 of the German Research Foundation (Deutsche Forschungsgemeinschaft, DFG) to B.W. funding E.P., a grant of the Leibniz-DAAD Research Fellowship Programme by the Leibniz Association and the German Academic Exchange Service (DAAD) to A.C.M. and N.D., and Ph.D. fellowships of the Landesgraduiertenförderung Sachsen-Anhalt awarded to C.N. and F.F. Financial support came from the Leibniz Association, the state of Saxony Anhalt, the Deutsche Forschungsgemeinschaft (DFG) Graduate Training Center GRK1026 “*Conformational Transitions in Macromolecular Interactions*” at Halle, and the Leibniz Institute of Plant Biochemistry (IPB) at Halle, Germany. To complete work on this project, a Short Term Scientific Mission (STSM) of the European Cooperation in Science and Technology (COST, www.cost.eu) was granted to A.C.M. and N.D. by the COST Action BM1307 – “*European network to integrate research on intracellular proteolysis pathways in health and disease (PROTEOSTASIS)*”. This work was partially funded by the grant DI 1794/3-1 of the German Research Foundation to N.D.

## AUTHOR CONTRIBUTION

A.C.M. performed the ubiquitination reactions and related analysis. E.P. and B.W. designed and synthesized the fluorescent probe, B.W. supervised the chemical synthesis, M.K. established PRT1 ubiquitination reactions, C.N. cloned and purified PRT1, F.F. cloned the X-eK-HAT fragment and performed site-directed mutagenesis. N.D. and A.C.M. designed the study, wrote the manuscript under consultation with all co-authors and designed the figures. All authors read and approved the final version of this manuscript.

## SUPPORTING INFORMATION

Additional supporting information may be found in the online version of this article.

## SUPPORTING FIGURES

**Supporting Information Figure 1. Modeled structure of the F-eK-Flv substrate and PRT1 N-terminal specificity.**

## SUPPORTING TABLES

**Supporting Information Table 1. State-of-the-art ubiquitination detection methods. Supporting Information Table 2. Oligonucleotides used in this study.**

## SUPPORTING METHODS

**Synthesis of the chemical probe NBD-NH-PEG_2_-NH-haloacetamide.**

## SUPPORTING REFERENCES

